# Considerations for the use of laboratory-based and field-based estimates of environmental tolerance in water management decisions for an endangered salmonid

**DOI:** 10.1101/2023.08.23.554483

**Authors:** Steven C. Zeug, Alex Constandache, Bradley Cavallo, Cramer Fish Sciences

**Affiliations:** Modeling Analysis and Synthesis Laboratory, 13300 New Airport Road, Suite 103, Auburn, CA 95602

## Abstract

Water infrastructure development and operation provides essential functions for human economic activity, health, and safety, yet this infrastructure can impact native fish populations resulting in legal protections that can, in turn, alter operations. Conflict over water allocation for ecological function and human use has come to the forefront at Shasta Reservoir, the largest water storage facility in California, USA. Shasta Reservoir supports irrigation for a multibillion-dollar agricultural industry, provides water for urban and domestic use, provides flood protection for downstream communities, and power generation as part of the larger Central Valley Project in California. Additionally, an endangered run of Chinook Salmon relies on cold water management at the dam for successful spawning and egg incubation. Tradeoffs between these uses can be explored through application of models that assess biological outcomes associated with flow and temperature management scenarios. However, the utility of models for management decisions are contingent on their characteristics, data used to construct them, and data collected to evaluate their predictions. We evaluated laboratory and field data currently available to parameterize temperature-egg survival models for winter run Chinook Salmon that are used to inform Shasta Dam operations. Models based on both laboratory and field data types had poor predictive performance which limits their value for management decisions. The sources of uncertainty that led to poor performance were different for each data type (field or laboratory) but were rooted in the fact that neither data set was collected with the intention to be used in a predictive model. Our findings suggest that if a predictive model is desired to evaluate operational tradeoffs, data must be collected for the specific variables desired, over an appropriate range of values, and at sufficient frequency to achieve the needed level of precision to address the modeling objective.

## Introduction

Development of water resources for human benefit is nearly ubiquitous in North American rivers. Modification of lotic ecosystems such as diversion for irrigation, urban use, hydropower development, and flood control has created significant economic value but is also associated with declines of native aquatic species that has led to some being listed under federal and/or state law (1,2,3). Once development of water resources has occurred, management and recovery efforts for legally protected species are largely constrained to occur in conjunction with continued human uses, and human uses must in turn consider the needs of protected species. The consequences of resource management decisions are magnified when these threatened or endangered species occur in the affected area. When water availability is limited, science can help inform allocation of resources through the application of models (4). However, the effectiveness of models to predict outcomes is closely tied to the characteristics of the data used to develop, parameterize, and apply the models.

There are two primary approaches to acquiring data on species’ responses to environmental drivers for use in models: (1) observational studies in the field; and (2) controlled laboratory experiments. There are tradeoffs associated with each approach that should be considered when using data from these methods to set management targets and predict outcomes. Field studies have the advantage of taking place in the habitats where the species of interest occurs, with elements of actual habitat structure and environmental conditions that can be difficult to replicate in the laboratory. However, under field conditions there are many variables simultaneously influencing outcomes that cannot be controlled. Thus, it can be difficult to confirm causal relationships between the species of interest and any single variable. Additionally, the range of conditions observed in the field is limited to the length of the study and the conditions that occurred during that time.

In contrast, laboratory studies have the advantage of controlling variables other than the one of interest and allowing for the investigator to set the range and magnitude of the variable being examined. Additionally, trials can be replicated to acquire sufficient statistical power to detect differences. The primary disadvantage of laboratory studies is that the experiments are often simplified and may not represent an appropriate range of field conditions and multivariable interactions, confounding the transfer of laboratory results to real-world conditions.

For poikilotherms like Chinook Salmon, temperature has a major influence on physiological performance. However, much remains unknown about the specific mechanisms of temperature limitation at different levels of biological organization (5) and environmental experience (6,7). Biological responses to temperature that are quantified under controlled laboratory conditions have been evaluated in the field with studies carefully designed for that purpose (8,9,10).

Differences between field and laboratory derived data have become prominent in water and temperature management decisions for Sacramento River winter run Chinook Salmon (*Oncorhynchus tshawytscha*), which are listed as endangered under the federal Endangered Species Act. Winter run are unique among California Chinook Salmon in that they spawn in the late spring and summer when meteorological conditions typically result in warm air and water temperatures. Historically, before the construction of Shasta Dam, winter run spawned in higher elevation habitat where water temperatures for spawning and egg incubation were more consistently cold during summer. Currently, there is a single population of winter run that spawns in the mainstem Sacramento River below Keswick Dam (the re-regulating reservoir for Shasta Dam) on the Sacramento Valley floor. Water temperatures in the downstream spawning areas during egg incubation are maintained using selective withdrawal from different water depths at Shasta Dam. Thus, management of cold-water volume in the reservoir, and release of that water into the river to maintain appropriate temperatures during the incubation period are critical components to winter run Chinook Salmon reproductive success.

Shasta Reservoir and associated facilities are also a critical part of the US Bureau of Reclamation’s Central Valley Project (CVP). Shasta is the largest reservoir in California and a key part of the CVP that provides water for roughly a third of California’s multi-billion-dollar agricultural industry and approximately one million households, generates hydroelectric power at both Shasta Dam and Keswick Dam, and provides flood control for the largest river in the state. These human needs must be considered in planning operations that affect cold water in Shasta Reservoir. There is considerable interest from resources managers to be able to predict winter run Chinook Salmon egg survival outcomes under different operational scenarios at monthly, annual and multi-year scales. Balancing human and ecological needs requires reliable information on species needs as well as effective monitoring of the life stages that can be affected by management decisions. As climate change continues to produce periods of extreme water scarcity and abundance, a framework for data collection for use in predictive models will be increasingly important for effectively managing water and biological resources.

Here we review the data available from laboratory and field studies to estimate temperature effects on survival of winter run Chinook Salmon during egg incubation. These are the primary data sources used in biological models to inform decisions on cold water management. This paper characterizes uncertainty in each data source, reviews assumptions needed to make predictions with models using these data and makes recommendations for studies and monitoring that can resolve conflicting conclusions drawn from each type of data.

## Methods

### Study site

The Sacramento River is the largest river in California, USA and drains a ∼ 70,000 km^2^ catchment including portions of the southern Cascade Mountains, Sierra Nevada Mountains and Coast Range. The river is impounded by Shasta Dam, completed in 1945, creating a reservoir with a capacity of ∼5.5 km^3^. Keswick Dam, located ∼15 km downstream of Shasta Dam serves as a re-regulating facility and is the current upstream limit for anadromous fish migration in the Sacramento River. The Mediterranean climate of California is strongly seasonal with cool wet winters and hot dry summers. This climate pattern drives variation in flow into the reservoir. However, flows below Shasta Dam have been significantly altered from pre-dam patterns with attenuation of peak flows in the wet season and higher flows in the dry season when agricultural and urban demands are greatest (11). To better manage cold water releases into the river, a temperature control device (TCD) was installed on Shasta Dam and became operational in 1997. The TCD can withdraw water from different depths to more efficiently access and blend water of different temperatures while continuing to generate hydropower. Below Keswick Dam the river flows south for 486 km before joining the San Joaquin River in the tidal Sacramento-San Joaquin Delta and then entering San Francisco Bay.

### Study species

Winter run Chinook Salmon are one of four runs that spawn in the Sacramento River, and are unique in that they enter freshwater in the winter while still immature and hold for weeks to months before spawning between May and July (the warmest and driest period of the year). Prior to the construction of Shasta Dam, winter run adults migrated to higher elevation habitat in the Sacramento, Pit and McCloud Rivers where temperatures were appropriate for egg incubation. Currently, the Sacramento River below Keswick Dam is the only location where winter run spawn, with most spawning in the first 9 kilometers where releases from Shasta Reservoir keep water temperatures cool. Winter run Chinook Salmon have been listed as endangered under the federal Endangered Species Act since 1994 and their population is supported by the Livingston Stone National Fish Hatchery that began operation in 1997.

### Laboratory data

Survival of winter run Chinook Salmon eggs as a function of temperature was evaluated in a laboratory study by the United States Fish and Wildlife Service (12). This study was a blocked design with five treatment levels corresponding to temperatures of 13.3, 14.4, 15.6, 16.7, and 17.8 °C and three replicates per treatment. Separate incubation units were used for each treatment level and individual replicates were placed in incubation cups containing 80-90 eggs each. The number of eggs for this study was limited by low abundance of this endangered species and all experimental eggs came from 3 spawning pairs. There were additional treatments where water temperatures were raised during incubation. Here, we focus only on the constant temperature portion of the study. Survival was assessed at four developmental stages following the accumulation of a specific number of accumulated temperature units (ATU) including, cleavage eggs (0-450 ATU), embryos (451-900 ATU), eleutheroembryos (901-1350 ATU), and pre-emergent alevins (1351-1800 ATU). Chillers and heaters were used to maintain target temperatures in each incubation unit. However, mean daily temperatures fluctuated up to ∼ 0.6 °C and individual measurements fluctuated up to 4.2 °C.

### Field data

Survival between egg deposition on the spawning grounds in the Sacramento River and migration of juveniles past a trap located approximately 95 km downstream of Keswick Dam is estimated as,

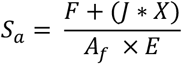

Where *S*_*a*_ is the annual estimate of egg-to-fry survival, *F* is the estimated number of juveniles migrating past a trap station at the fry stage, *J* is the estimated number of juveniles migrating past the trap station at the pre-smolt and smolt stage, *X* is the inverse of an estimate of the survival rate between the fry and pre-smolt/smolt stage upstream of the trap site, *A*_*f*_ is the estimated number of adult females and *E* is an estimate of mean fecundity for adult females. The estimates needed for this calculation have been available since 1996 except for two years (2000 and 2001) when no juvenile trapping occurred.

The number of females is estimated by the California Department of Fish and Wildlife with a mark-recapture survey of spawner carcasses and a Cormack-Jolly-Seber model. Additionally, the distribution of redds above and below the survey reach and data from fish taken into Livingston Stone National Fish Hatchery are used to inform the estimate of female spawners. Detailed methods for the carcass survey and estimation procedures can be found in (13).

The number of fry is estimated at a sampling station operated by the United States Fish and Wildlife Service located approximately 95 km downstream of Keswick Dam near the city of Red Bluff. At this station, three or four rotary screw traps are deployed to capture juvenile salmonids as they migrate downstream. Mark-recapture trials using fish captured in the traps are used to estimate trap efficiency and expand trap counts to total passage. Multiple runs of Chinook Salmon spawn upstream of the juvenile trapping station and the run of fish captured is determined using a length-at-date model that uses assumptions about spawn timing and growth to separate the runs by size on a specific date. Additionally, juveniles express multiple migration strategies with some fraction passing the trap as fry soon after emergence and others rearing upstream of the trap for weeks or months and then migrating as pre-smolts or smolts. For the egg-to-fry survival calculation, pre-smolts and smolts are converted to “fry equivalents”. This is done by assuming the survival rate between the fry stage and pre-smolt/smolt stages is 59%. This value was obtained from an undated paper by Richard J. Hallock of the California Department of Fish and Wildlife, that compared mean adult return rates from fish released as yearlings relative to young of the year smolts.

Sub-daily temperature data is monitored at three stations within the spawning reach, although the data from the entire period is not available at all stations. The River Assessment and Forecasting Tool (RAFT) is a model that estimates surface water temperature at 1km intervals and has been used to assess temperature conditions in the spawning reach (14).

The time and location of egg deposition in the study area can be estimated in two ways. The first approach uses data from a mark-recapture survey of winter run carcasses performed by the California Department of Fish and Wildlife discussed above. Surveys of the spawning area are performed by boat multiple times per-week during the season (> 40 surveys) and the distribution of female carcasses from this survey can be used to assess the spatial and temporal pattern of egg deposition. Under the second approach, aerial surveys are conducted during the spawning season to visually identify new redds that have been constructed between surveys. Each year, 7-16 survey flights are conducted within the spawning season to visually identify new redds constructed between flights. The exact number of flights can be affected by aircraft availability and project funding. Additionally, water clarity and depth limit the ability to identify redds with this method.

### Temperature-mortality models

The USFWS (12) conducted analysis of variance to assess differences among treatments but did not create a predictive model of egg mortality as a function of temperature. A predictive model of egg mortality as a function of temperature based on the USFWS data was constructed by Zeug et al. (15) as part of a lifecycle model for winter run Chinook Salmon. In that effort, the mean survival value for each temperature treatment through the pre-emergent alevin stage was fit using an exponential function.

To incorporate the variation within treatments, the USFWS data were refit using individual replicates within each of the temperature treatments as individual observations rather than treatment means. Additionally, we calculated survival through the eleutheroembryonic stage. This was done because once the pre-emergent alevin stage is reached, fish can ventilate their gills and have some mobility within the gravel. This would allow them to respond to stressors differently than when confined within the egg chorion at the location of deposition. Similar to Zeug et al. (15), we fit an exponential model to these individual observations and calculated confidence and prediction intervals for the relationship.

An analysis of the field data was performed by Martin et al. (16) with non-linear regression to estimate the temperature below which no mortality occurs (T_c_), the rate of mortality after exceeding T_c_ (β_t_), the background survival rate (S_0_) and the carrying capacity of female spawners (K). Although all the field data are estimates, Martin et al. (16) treat them as observations without error. We reanalyzed those same data, plus an additional 6 years, with Bayesian rather than frequentist methodology to fit each parameter.

We analyzed twenty-four years of field data (1996-1999, 2002-2021) with a faithful Bayesian translation of the frequentist model used by Martin et al. (16). Temperature measurements were provided as an R by C matrix, where R is the total number of observed redds, and C is the maximum number of temperature available for a redd. Some redds had less than C measurements, in which case the extra entries in their row were filled with NaN (not a number) values. An R dimensional vector associates each redd with the year in which it was observed, while a Y dimensional vector provides the observed overall ETF survival rate for each of the Y years for which observations were available.

For every redd *r = 1, 2, …, R*, the thermal hazard is assumed to follow the Martin et al. (16) formula:

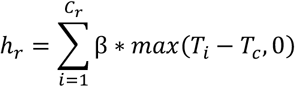

The overall survival rate is then given by the formula:

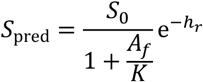

Where *K* is a carrying capacity and *S*_*0*_ is the background survival rate. The observed ETF mortality is assumed to be normally distributed around the theoretical value, on a logit scale:

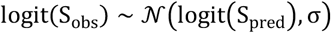

where logit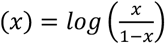

## Results

### Lab model

There was considerable variation in survival within each temperature treatment with the exception of the 17.8 C treatment where all eggs died prior to emergence. In the lowest temperature treatment (13.3 C) mean survival was 83.3% and ranged from 68.9%-95.7%. In the 14.4 C treatment survival was 69.5% (range 49.4%-94.4%), the 15.6 C treatment had a mean survival of 21.9% (range 6.9%-41.6%) and at 16.7 mean survival was 5% (range 0% -11.6%).

An exponential model fit the daily mortality values from the laboratory data well (F_1,13_ = 57.17, p < 0.001) and explained 81.5 % of variation (Figure 1). The 95% confidence interval for the relationship was relatively wide reflecting the high variation within treatments (Table 1). Prediction intervals were also wide, suggesting relatively low precision for predicting future survival rates throughout the range of temperatures examined.

**Table 1.** Mean predictions from the exponential model fitted to the laboratory data with confidence intervals and prediction intervals for each treatment level.

| Temperature | Model predicted survival | Confidence interval | Prediction interval |
| --- | --- | --- | --- |
| 13.3 | 88.2 | (76.1, 94.4) | (43.4, 98.1) |
| 13.9 | 79.9 | (65.0, 88.9) | (23.9, 96.5) |
| 14.4 | 67.1 | (50.3, 79.3) | (8.4, 93.7) |
| 15 | 49.5 | (32.4, 64.5) | (1.3, 88.9) |
| 15.6 | 29.2 | (14.5, 45.6) | (0.0, 81.4) |
| 16.1 | 11.8 | (3.1, 26.7) | (0.0, 70.3) |
| 16.7 | 2.5 | (0.1, 12.1) | (0.0, 55.3) |
| 17.2 | 0.2 | (0.0, 3.8) | (0.0, 37.8) |
| 17.8 | < 0.1 | (0.0, 0.7) | (0.0, 20.7) |

**Figure 1.**
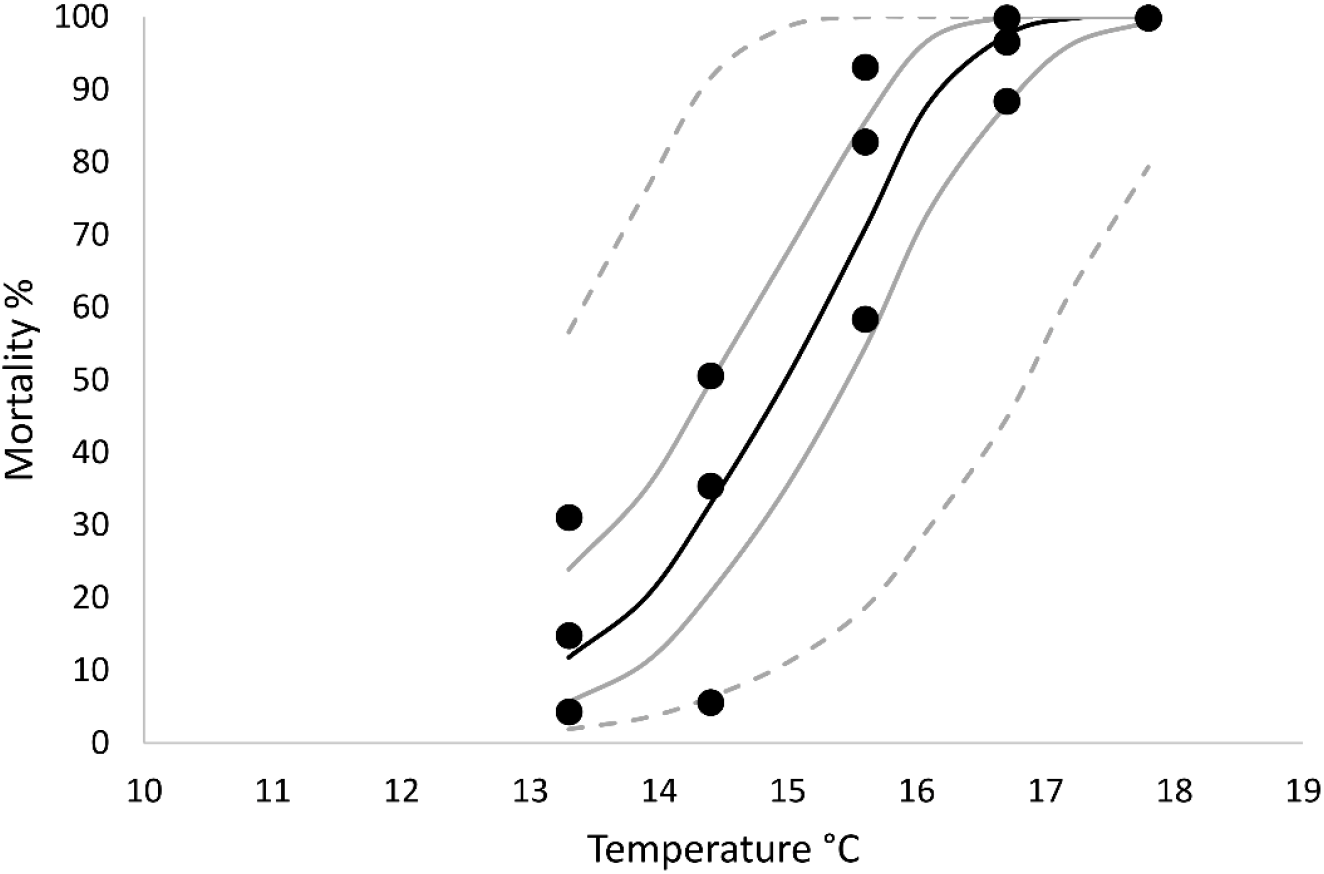
Exponential model fit to mortality rates from USFWS 1999. Observed values are represented by filled circles, the solid black line is the mean model estimate, the solid gray line is the 95% confidence interval, and the dashed gray line is the 95% prediction interval.

### Field model

The distribution of parameter values produced by the Bayesian analysis fell along a curve rather than a single point indicating that the model parameters were not identifiable (Figure 2). A similar curve is apparent in Figure 3 of the publication by Martin et al. (16) that analyzed a subset of these data (excluding years 2016-2021) using frequentist methodology.

**Figure 2.**
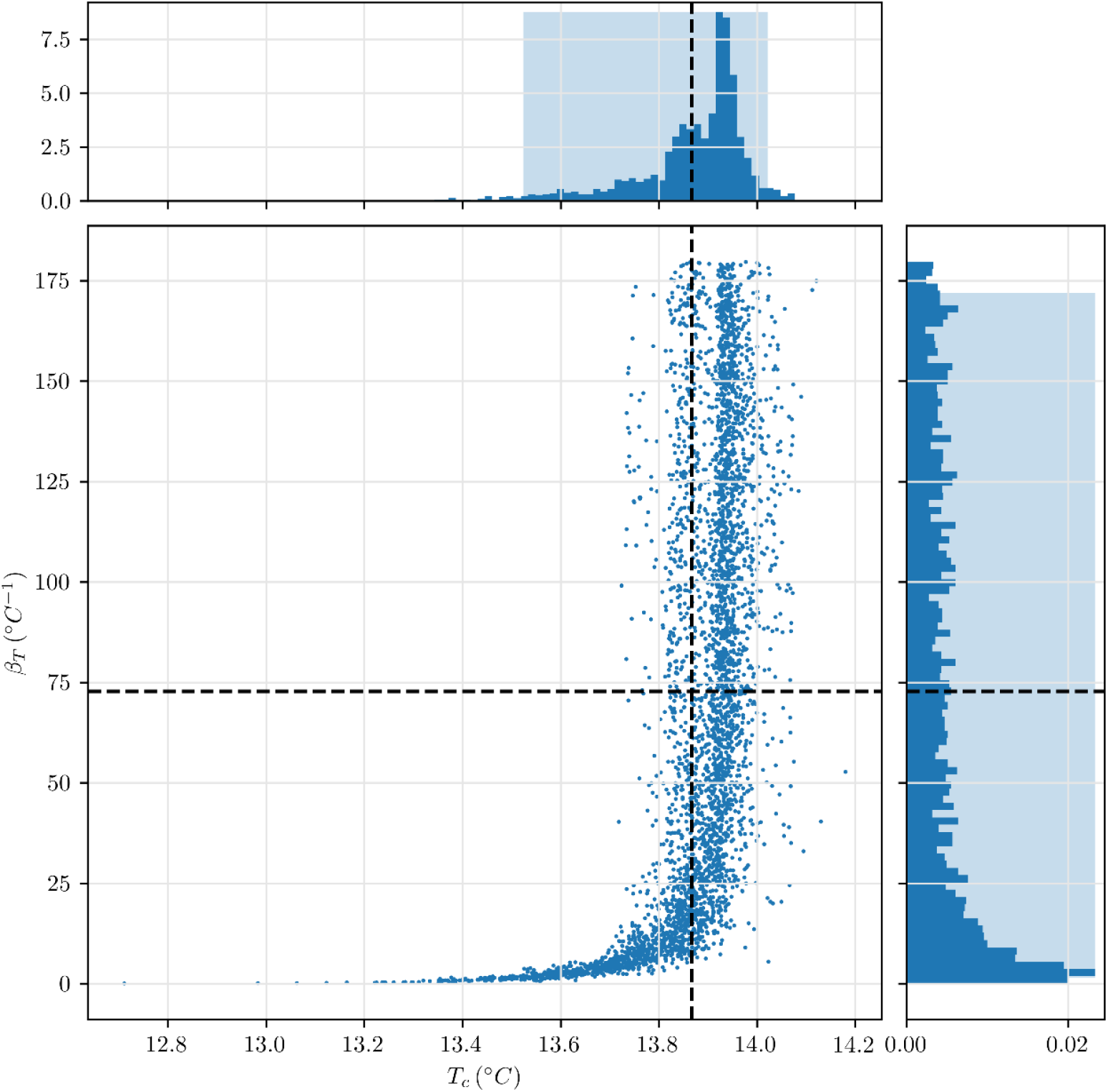
Biplot of estimates for *T*_*c*_ and *β*_*t*_ from Bayesian fitting of field data assuming fixed values of background survival (*S*_*0*_ = 0.36) and carrying capacity (*K* = 9,107).

**Figure 3.**
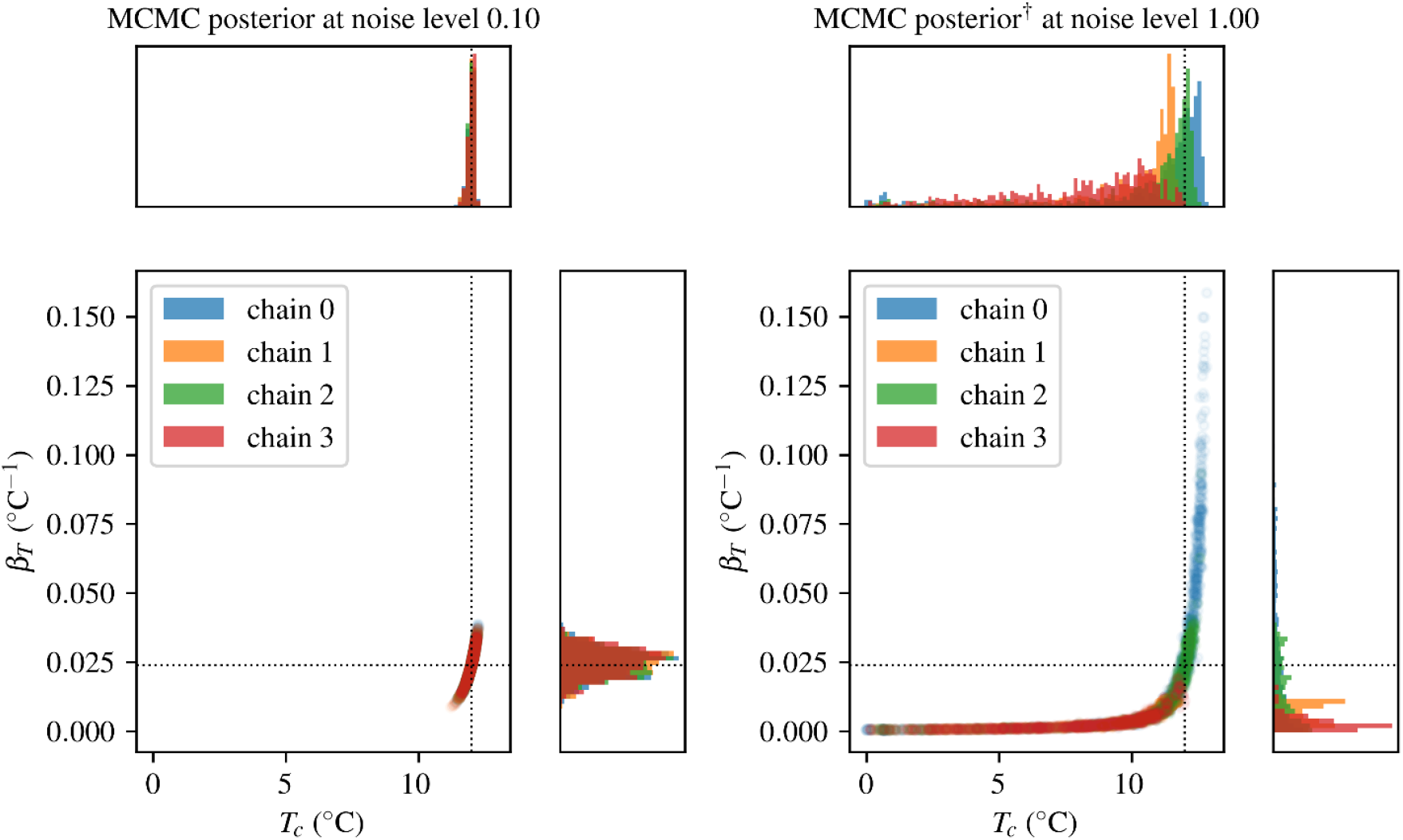
Outputs from the analysis of synthetic field data with known values of *T*_*c*_ (12.0) and *β*_*t*_ (0.024). Panel A are the results with minimal noise introduced into the data (α = 0.10) and panel B are the result with a more realistic level of noise in the data (α = 1.0).

To further explore the potential of using the available field data within this model framework to obtain identifiable parameters we employed a simulation approach (17). We generated synthetic data to approximate real observations that would occur where *T*_*c*_ = 12.0°C and *β*_*t*_ = 0.024 °C^-1^*d*^-1^—the values for the hazard model parameters claimed as “best fit” in Martin et al (16). Background survival (S0) was fixed at 36% and carrying capacity (K) fixed at 9,107. These values were also the best fit values from Martin et al. (16). With minimal noise introduced into the synthetic observations (α = 0.1) the Metropolis-Hastings algorithm yields tight, unimodal posterior distributions for *T*_*c*_ and *β*_*t*_ centered at the “best fit” values in Martin et al (16) (Figure 3). However, at a more realistic noise level (α = 1.0) different chains of the Metropolis-Hastings algorithm concentrate on different portions of the curve, producing different posterior distributions and maximum likelihood estimates differing significantly from the “best fit” values (Figure 3).

### Summary of results methods

The laboratory data indicate a strong exponential relationship between egg survival and temperatures above 13.3 °C. However, this relationship suffers from high uncertainty which makes management outcomes difficult to predict.

The field data currently used to monitor egg-to-fry survival includes multiple life stages (adults, eggs, fry, smolts) that have different sensitivities to temperature and are exposed to various other sources of mortality. Thus, the field monitoring data cannot be reliably used to validate predictions of temperature-dependent effects on eggs produced by the laboratory-derived model. The field-based model attempts to account for non-temperature mortality sources in the monitoring data to estimate a critical thermal maximum (*T*_*c*_) for eggs and rate of mortality (*β*_*t*_) but assumes that all mortality sources for each life stage are constant among years with the exception of temperature and density dependent effects on eggs. The reliability of the parameter estimates are contingent on the acceptance of that assumption. However, the lack of model identifiability suggests that reliable parameters cannot be estimated due to the nature of the monitoring data and model structure.

## Discussion

Water management decisions at Shasta Dam can directly affect the most endangered run of Chinook Salmon in the Sacramento River during a sensitive life stage. These decisions also affect California’s multi-billion-dollar agricultural industry, power generation, and the water supply for one million households. Competition between multiple uses results in trade-offs that must be assessed by water managers to determine the timing and volume of water allocations. Predictive models of biological responses to physical drivers can be a powerful tool to improve operational decisions and provide acceptable outcomes; particularly when resources become limiting. However, the effectiveness of these models is limited by the characteristics of the data used to develop, parameterize, and apply the models, and evaluate their predictions. Our evaluation of the laboratory data and field-derived data used to parameterize temperature-egg survival models suggested that both sources have significant uncertainty that constrains their utility for management. This finding became apparent most recently during the third year of drought in California (2022) when water resources were limited, and the field-based model proposed by Martin et al. (16) was used to assess management options. Despite more favorable temperatures than expected, the egg to fry survival monitoring in the field was the lowest on record; far below the value predicted by the model (18). Thiamine deficiency was detected in winter run spawners which can cause high mortality of hatched fry. Thus, poorly understood sources of mortality likely contributed to poor predictive performance. Given the enormous economic and natural resource value of water stored behind Shasta Dam, better data are needed to parameterize models with sufficient performance for predicting survival and evaluating management trade-offs.

Models describing temperature effects on survival of winter run Chinook Salmon eggs were uncertain regardless of the data source/model used which complicates attempts to accurately predict the outcome of water management decisions for Shasta Dam. The model parameterized with the laboratory data produced reasonable confidence intervals; however, when models are used to predict future outcomes, prediction intervals are more appropriate to characterize uncertainty (19). In the laboratory data, variation in survival within treatment levels resulted in prediction intervals sufficiently wide that there would be low confidence in survival outcomes from managing to a specific water temperature. The model parameterized with field data was not identifiable and reliable coefficient values could not be estimated. Therefore, the field data model does not appear to provide a reliable basis for evaluating operational tradeoffs. Direnzo et al. (17) described that non identifiability frequently happens when two or more variables “trade off” to produce the same outcome with equal likelihood. In this case, *T*_*c*_ and *β*_*t*_ trade off coefficient values where eggs can die rapidly at higher temperatures (higher *T*_*c*_ and higher *β*_*t*_) or slowly at lower temperatures (lower *T*_*c*_ and lower *β*_*t*_). Further, estimates of carrying capacity (*K*) and background survival (S_0_) interact with *T*_*c*_ and *β*_*t*_ to allow more equally likely combinations to produce the same value of survival. Without strong prior information for at least one parameter from an independent source, a wide range of coefficient values for each parameter are equally likely (20).

In both the field and laboratory studies, the data used to model survival effects were not collected for that specific purpose and that is reflected in the predictive models. The laboratory data were collected to evaluate early-life stage survival under different, but constant, temperature regimes rather than to identify a specific threshold temperature when temperature dependent mortality effects commenced. The low number of available eggs for endangered winter-run Chinook Salmon resulted in a small number of replicates for each treatment level (n= 3). The study found significantly higher cumulative mortality at temperatures above 13.3 C. However, the small number of replicates indicates there was likely to be insufficient statistical power to detect differences even if they did exist (Type II error). This uncertainty resulted in wide prediction intervals of the exponential model.

The field data were not collected to estimate egg survival but rather are individual components of multiple monitoring efforts each with their own, different objectives, including carcass surveys to estimate escapement, redd surveys to estimate spawn timing and spatial distribution, emigration monitoring to estimate juvenile migration timing and abundance, and fecundity monitoring of brood stock to estimate hatchery production. Estimating egg survival from currently available field data requires five separate model inputs each with an associated uncertainty. However, those uncertainties are not incorporated by the Martin et al. (16) modeling effort or the Bayesian analysis presented here. The simulation analysis indicates that with realistic levels of noise in the data, known values of model parameters cannot be confidently recovered.

The importance of appropriate data and incorporation of uncertainty for predictive modeling in ecology is well recognized (21, 22).Given the endangered status of winter run Chinook Salmon and the importance of Shasta Dam and reservoir operations for anthropogenic uses, along with the known importance of temperature to survival of Chinook Salmon eggs, it is important to obtain data that can provide better predictions for use in water management decisions. The potential remedy to reduce uncertainty differs for each approach. For laboratory studies, primary considerations would be to improve data quality for modeling by increasing replicates at each treatment level. Given that winter run Chinook Salmon are endangered, and adaptation to local conditions is known to occur in different populations of Chinook Salmon (7,23), studies may need to occur in years when sufficient numbers of adult winter run are available. Additionally, care is needed to account for parentage in each treatment level as this can have a substantial effect on survival outcomes regardless of treatment effects (24,25). Finally, experimental set ups should be designed to replicate natural conditions as much as possible to increase confidence in transferring results to conditions *in situ*. This may include the use of gravel substrates in experimental units, providing representative intergravel flow rates, inducing diel temperature fluctuations, and managing other variables (e.g., dissolved oxygen) to mimic natural patterns.

Field studies can be improved by using methods and study designs that focus directly on the stage of interest. Survival of salmonid embryos in the field has been a topic of considerable research, and is associated with a well-understood, accepted methodology (e.g. 25, 26, 27). Appropriate methods for estimating egg-to-fry survival in the field include burial of a known number of eggs in various designs of incubation containers so fry can be captured and directly counted at hatch, or when they emerge from the gravel. Remote monitoring devices and loggers can be buried along with experimental egg boxes to obtain data on the exact temperature and other conditions experienced during incubation. Temperature-embryo development models are used to determine when excavation of experimental units should occur and data on substrate size can be evaluated at that time. Careful attention to field study design is necessary to avoid confounding effects of uncontrolled variables while maintaining desired treatment effects. For example, Merz et al. (28) found that there was an unexpected interaction between substrate size and egg predator abundance in an evaluation of substrate size effects on egg survival. Thus, even when the experimental design is sound, unexpected interactions can occur in field settings that introduce mortality not attributable to treatment effects.

Management decisions around Shasta Dam operations have significant biological and economic consequences. However, our evaluation of existing data sources and models suggest that the ability of current models to predict winter run Chinook Salmon egg survival outcomes from water temperature is poor. Model uncertainty can come from multiple sources including model conceptualization, model development, data development and parameter estimation. For laboratory and field studies, none of the data relied on were collected for the purpose of developing predictive models of temperature effects on eggs. Therefore model development and conceptualization was constrained by the type of data available. Increasing model predictive performance requires data collected using a study designed with that intended purpose. Laboratory studies have been the gold standard in science because of the ability to control confounding variables. However, if a laboratory-based model is to be used for predicting outcomes in the field, those confounding variables and their interactions need to be considered. Regardless of data source, validation of models with direct observation of egg survival outcomes under field conditions will be an essential metric to evaluate the value of predictive models.

## Acknowledgements

We thank the National Marine Fisheries Service Southwest Fisheries Science Center for providing the code and data for the frequentist field model. Lee Bergfeld and Mike Deas provided valuable comments on previous drafts of the manuscript. Funding for this effort was provided by the Sacramento River Settlement Contractors.

## Notes

https://github.com/fishsciences/wr_etf_data.git

### Competing Interest Statement

The authors have declared no competing interest.

