## Supplement 1 for "Considerations for the use of laboratory-based and field-based estimates of environmental tolerance in water management decisions for an endangered salmonid"

### What is (non-)identifiability?

A model with parameter(s) $\theta$ is said to be non-identifiable if two or more distinct parameter values are observationally equivalent. In mathematical terms, the probabiliy distributions $p(d|\theta_1)$ and $p(d|\theta_2)$ are equal, where $d$ is any set of observations and $\theta_{1,2}$ are two distinct sets of parameters. In data analysis practice, a weaker version of non-identifiability is used: the model is considered non-identifiable if two (or more) significantly different parameter choices $\theta_1$ and $\theta_2$ produce nearly identical observation probability distributions $p(d|\theta_1) \approx p(d|\theta_2)$.

There are, broadly speaking, two kinds of non-identifiability: structural and practical. Structural non-identifiability arises when the model depends on the parameters through a non-injective function $f$:

$$p(d|\theta) \equiv p\big(d|f(\theta)\big)$$

In this case, the level sets of $f$, namely $\{\theta \mid f(\theta) = a\}$ are observationally equivalent, as they produce the same probability distribution over observations.

Here is a simple example of structural non-identifiability: suppose we have a factory that produces gadgets. With some non-zero probability, a gadget coming off the assembly line may be defective. The defects can be of two types: type A are defects introduced in our assembly process, while type B are defects caused by defective components sourced from our suppliers. If $p_A$ and $p_B$ are the probabilities of the two types of defects, then the probability that a deffective gadget is produced is:

$$p = p_A + p_B - p_A\, p_B$$

If we randomly tested gadgets coming off the assembly line we could precisely measure $p$ if the sample size was large enough, but increasing the sample size would not allow us to precisely infer the value of $p_A$, as there are infinitely many combinations of $p_A$ and $p_B$ that produce exactly the same value of $p$. By measuring $p$ we simply constrain the pair $(p_A, p_B)$ to lie on one of the curves in the following plot:

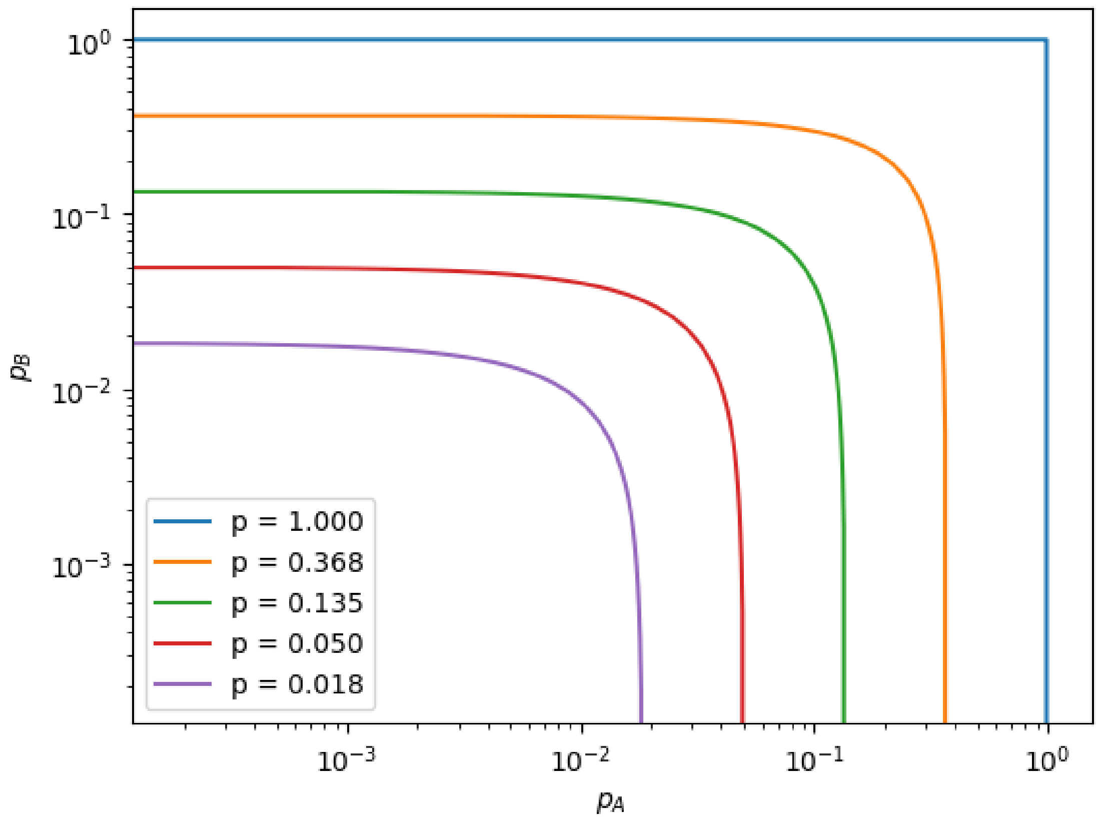


As we can see, for any value of $p$, the individual probabilities $p_A$ and $p_B$ can vary over multiple orders of magnitude. The Martin et al. (2017) approach to analysis of the field data has a similar problem: we measure overall mortality ($M$), which is the cummulative result of three independent types of mortality: background mortality ($M_0$), competition-related mortality (M_C), and temperature-related mortality ($M_T$). These quantities are only constrained by the formula:

$$1 - M = (1 - M_0)(1 - M_C)(1 - M_T)$$

Obviously, the same overall mortality M can be realized through infinitely many combinations of $M_0$, $M_C$ and $M_T$. For example, $M = 0.8$ can be achieved by setting $M_0 = 0$, $M_C = 0$, and $M_T = 0.8$, or by setting $M_0 = 0.8$, $M_C = 0$, and $M_T = 0$. These two allocations would result in very different estimates for $\beta_T$ (the rate at which eggs die) and $T_c$, (the temperature below which there is not temperature-dependent mortality),the two parameters used to model temperature-dependent mortality $M_T$ in Martin et al. In the latter case the parameter $T_c$ would have to be larger than any temperature observed in our dataset, while in the former we would expect that many of the temperature observations substantially exceed $T_c$.

This presents an insurmountable challenge when trying to fit the model to field data, where the three types of mortality cannot be disaggregated. This problem can only be circumvented by controlled measurements in the lab. In such a controlled setting there would be no competition between spawners, so we could assume that $M_C = 0$. A control group of eggs could be incubated at the known optimal incubation temperature, and their measured mortality would provide a direct measurement of $M_0$. Other egg baskets could be exposed to various high temperature profiles. With $M_0$ and $M_C$ known, we could calculate their $M_T$ value, which could then, in theory, be used to infer $\beta_T$ and $T_C$.

Even if we could disaggregate $M_T$ from the overall mortality $M$ in field measurements, fitting the model to field data may still present serious challenges, because the model is also afflicted with a second kind of non-identifiability: practical non-identifiability. This type of non-identifiability occurs when the sensitivity matrix of the model relative to the parameters becomes ill-conditioned (detail below), making it extremely difficult for any numerical inference algorithm, frequentist or Bayesian, to converge to the ground truth.

Suppose we have a model linking a dependent quantity $y$, to a vector of predictors $\mathrm{x} = (\mathrm{x}_1, \mathrm{x}_2, \ldots \mathrm{x}_\mathrm{n})$, and the model is parameterized by $\theta = (\theta_1, \theta_2, \ldots, \theta_k)$, that is to say $y = f(\mathbf{x}; \theta)$. If we wanted to fit this model to a set of observations, $(y_1, \mathbf{x}_1), (y_2, \mathbf{x}_2), \ldots, (y_N, \mathbf{x}_N)$, we would need to minimize the function

$$\chi^2(\theta) = \sum_{i=1}^{N} \left(y_i - f(\mathbf{x}_i; \theta)\right)^2$$

For this to be possible in practice, a necessary (but not sufficient) condition is that the sensitivity matrix

$$H(\theta) = \begin{pmatrix} \dfrac{\partial f(\mathbf{x}_1; \theta)}{\partial \theta_1} & \cdots & \dfrac{\partial f(\mathbf{x}_1; \theta)}{\partial \theta_k} \\ \vdots & & \vdots \\ \dfrac{\partial f(\mathbf{x}_N; \theta)}{\partial \theta_1} & \cdots & \dfrac{\partial f(\mathbf{x}_N; \theta)}{\partial \theta_k} \end{pmatrix}$$

must have linearly independent columns. If this holds, and we start the optimization from an initial guess that is located sufficiently close to the minimum, an optimization algorithm will always converge on the minimum. In practice, even when this condition holds, but the columns of $H(\theta)$ are very close to being linearly dependent, numerical algorithms will become so sensitive to rounding errors that they will converge to a random value every time we run them.

In the case of the Martin et al. (2017) model, $y$ is the hazard, $\mathbf{x}$ is the set of daily temperatures $(T_1, T_2, \ldots, T_n)$ for one redd, and $\theta = (\beta_T, T_c)$. The model function $f$ is:

$$f(T_i;\, \beta_T, T_c) = -\beta_T \sum_i \max(T_i - T_c, 0)$$

while the sensitivity matrix has two columns, corresponding to the two parameters, $\beta_T$ and $T_c$:

$$H(\beta_T, T_c) = \begin{pmatrix} \sum_{T_{1i}>T_c} (T_{1i} - T_c) & -\beta_T \sum_{T_{1i}>T_c} 1 \\ \vdots & \vdots \\ \sum_{T_{Ni}>T_c} (T_{Ni} - T_c) & -\beta_T \sum_{T_{Ni}>T_c} 1 \end{pmatrix}$$

Consider, for example the case when all the temperature measurements for a particular redd fall below $T_c$. In such a case all the terms under the sum in the formula for the hazard function vanish, for any value of $\beta_T$. In other word, this observation provides a constraint on $T_c$, namely $T_c > \max_i(T_i)$, but does not constrain $\beta_T$ at all. All possible values of $\beta_T$ are equally compatible with this observation. If all the redds in our dataset only contained temperature observations below $T_c$, then we could not expect to be able to constrain $\beta_T$ using such a flawed dataset. This can also be seen from the fact that all the entries in the sensitivity matrix vanish, making its columns linearly dependent.

If the dataset improved a bit, and a few redds had one or two temperature readings above $T_c$, then, by continuity arguments, the situation would improve only slightly and we would be able to restrict $\beta_T$ to a fairly wide interval, but not to a precise value

This situation might seem extreme, but in the actual dataset, more than 40% of redds don't have any temperature readings above the reported value of $T_c = 12^\circ$C, while overall less than 18% of all the temperature readings exceed the reported $T_c$, so the available data is closer to such a pathological example than it is to an ideal experiment design.

We can visualize the level curves of the sum of square errors function of the Martin et al. model for some synthetic data, generated using the following procedure.

- draw a $20 \times 80$ matrix of random temperatures from a normal ditribution with the same mean and variance as in the Martin et al. dataset ($\mu_T = 11.35\,^\circ$C, $\sigma_T = 0.75^\circ$C). These represents daily incubation temperatures for 20 redds.
- using the reported values for $T_c$ and $\beta_T$, compute the theoretical hazard value $H_0$ for each redd.
- add some lognormal$(0, \sigma_H)$-distributed noise to each $H_0$, to obtain the observed hazard $H$.
- at the nodes of a regular grid in the $(T_c, \beta_T)$ plane compute the sum of square errors for this synthetic dataset, using the formula:

$$\chi^2(\beta_T, T_c) = \sum_{i=1}^{20} \left( H_i - \beta_T \sum_{j=1}^{80} \max\left(T_ij - T_c, 0\right) \right)^2$$

First let's examine the sum of square error (SSE) landscape when the distribution of temperatures is as in the real dataset, and there is no observation noise ($\sigma_H = 0.$):

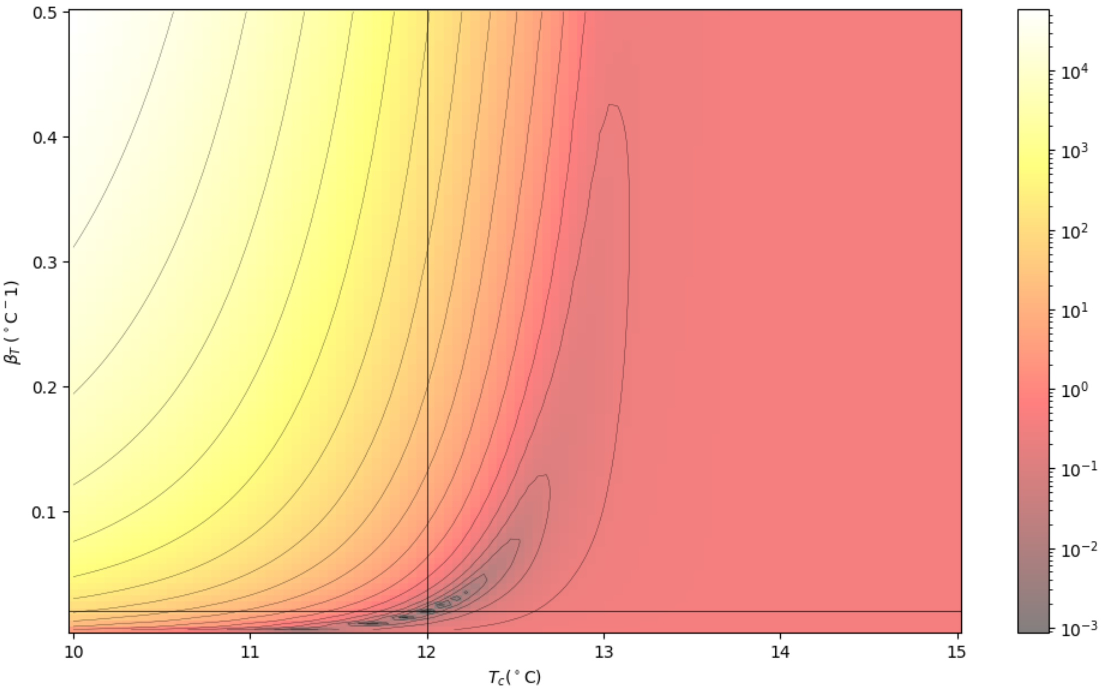


Even under such ideal conditions we can immediately see several signs of trouble. The first one is the banana-shaped level sets, a tell-tale sign of non-identifiability. Next is the fact that while the $\chi^2$ function has a local minimum at the ground truth (marked by the crosshairs), it also has other spurious local minima which develop because of numerical ill-conditioning. An optimization algorithm may converge to a different one of them every time it is run. These spurious local minima span more than 1 °C along the $T_c$ axis.

Next, let's keep the temperature distribution the same, but increase the observation noise. The individual minima coalesce into a banana-shaped valley with a flat bottom. The optimizer is free to settle anywhere therein. The valley spans several orders of magnitude along the $\beta_T$ axis.

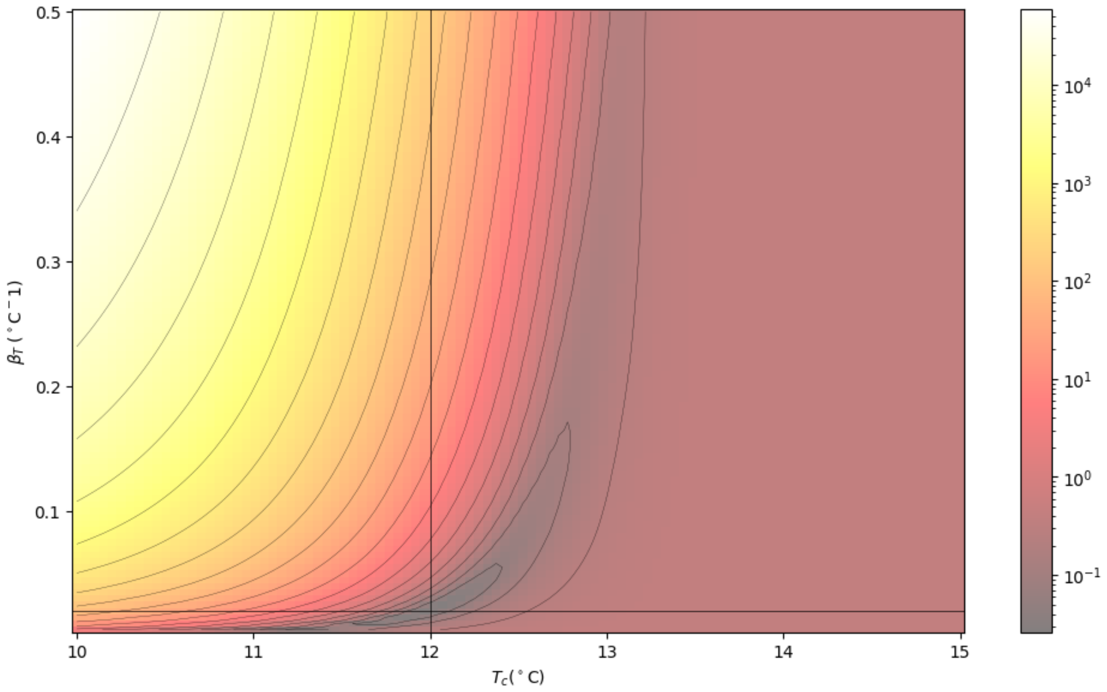


Next let's change the temperature distribution so that most readings fall below the true value of $T_c$. We will mark the minimum observed temperature in each redd by a blue vertical line, and the highest by a red one. The ground truth is no longer the most pronounced local minimum. Even at small noise levels, spurious local minima develop at very large values of $\beta_T$. It is now imposible to tell where the true value of $\beta_T$ might be located, just as we expected based on our previous qualitative analysis of a dataset where most observations fall below $T_c$:

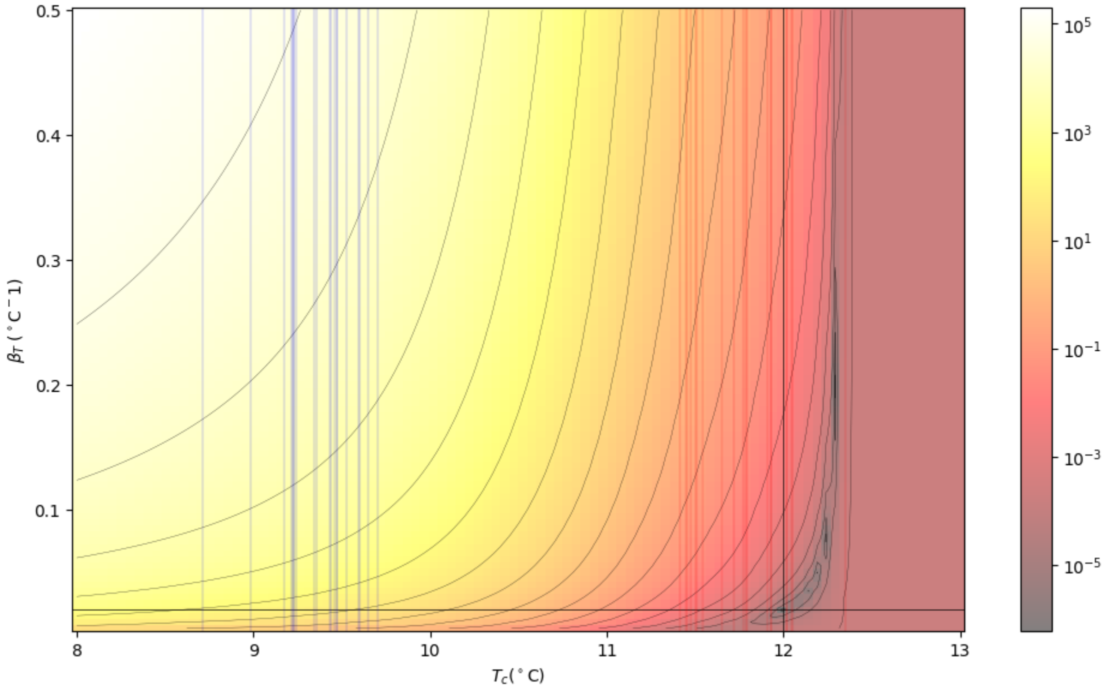

If we go in the opposite direction and generate a dataset where most temperature readings are above the true $T_c$, the spurious minima develop on the horizontal branch of the valley, rendering $T_c$ unidentifiable, even at low levels of noise:

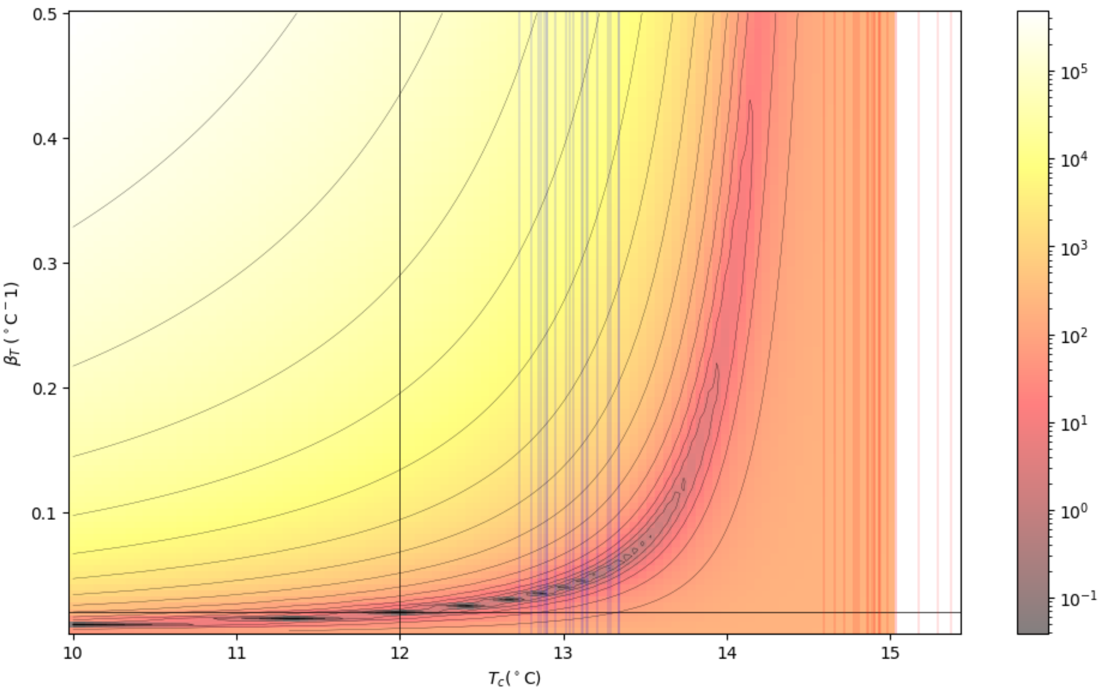


We see here that most of the spurious local minima are located in the elbow of the level sets, but the location of the elbow is constrained not by the ground truth, but by the distribution of observed temperatures. If we ran the optimizer a large number of times, it would settle most often in the elbow of the curves, leading us to wrongly conclude that the ground truth is located there.

To understand why these things are happening we need a way to visualize how close the sensitivity matrix is to linear dependence at various points in the $(\beta_T, T_c)$ plane. Since in our case the sensitivity only has two columns, a good measure of their linear dependence as vectors is the absolute value of the cosine of the angle between them. This quantity ranges from 0 to 1, being 1 when the two vectors are collinear (parallel or anti-parallel), and 0 when they are orthogonal. The system will be practically non-identifiable whenever this value is very close to 1.

If we plot this collinearity measure for different temperature distributions, we get the following plots:

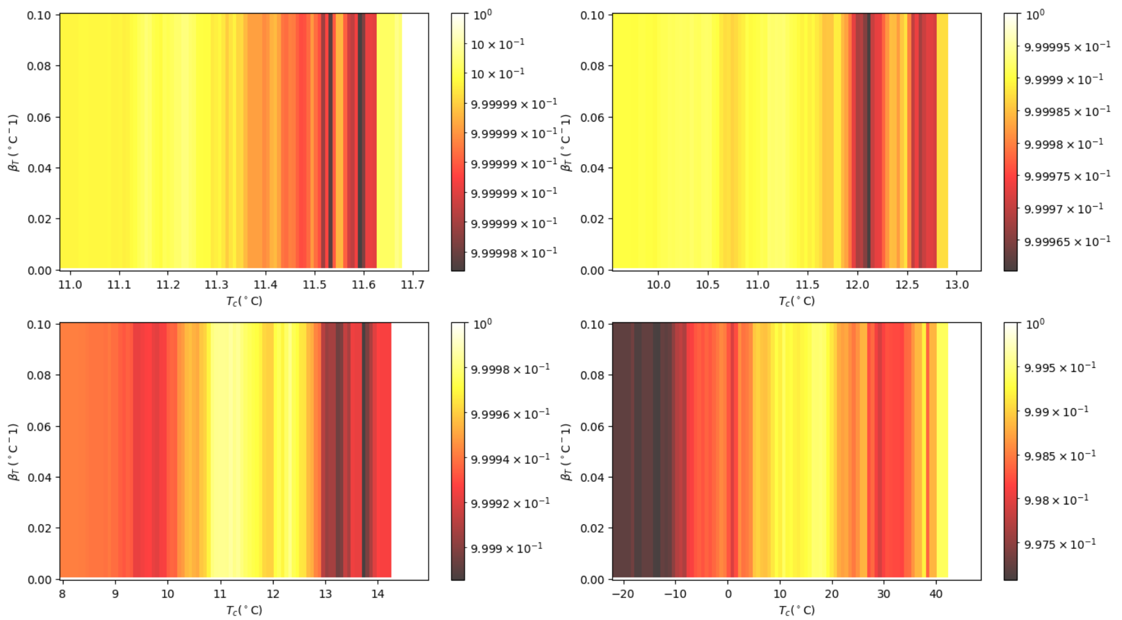


We see from these plots that unless the observed temperatures span an unrealistically large range of values, as in the lower right plot, the collinearity measure of the sensitivity matrix never goes below 0.999.

In conclusion, the Martin et al. model has an SSE function that presents elongated, banana-shaped level sets, which pose serious convergence challenges to numerical optimization algorithms. Multiple false local minima develop along the bottom of these regions, due to numerical instabilities, which make identification of the true values of the model parameters impossible. We could only hope to fit the model to a dataset if the following three conditions can be satisfied:

1. We can measure mortality very precisely.
2. We disaggregate the temperature related mortality.
3. The incubation temperature time series must be carefully selected to improve the conditioning of the sensitivity matrix and avoid numerical artifacts.

These conditions may only be satisfied in a lab setup, and even then identifiability would be problematic.
